# Microcalorimetry reveals bacterial motility as an expensive trait during starvation

**DOI:** 10.64898/2026.10.11.757999

**Authors:** Johannes M. Keegstra, Olivier Braissant, Riccardo Foffi, Blake Ushijima, Oliver Müller, Victoria J. Orphan, Roman Stocker

## Abstract

Quantifying the energetic costs of cellular traits is important to understand their evolutionary constraints and ecological function. However, direct measurements of energy expenditure of distinct traits are rare. Here, we present direct measurements of the power dissipation due to bacterial flagellar motility using microcalorimetry, based on measuring the power dissipation of a motile population and a non-motile mutant. For the single-flagellated marine bacterium *Vibrio coralliilyticus*, we determined the power dissipated by motility in cultures to be 0.91 µW/mL/OD. Per swimming cell this amounts to 2.0 ×10^*−*15^W, or a turnover of 2.5 ×10^4^ ATP molecules per second. The power dissipated for motility equals 35 % of total metabolism during starvation, indicating this is likely the most expensive cellular trait during starvation. The dissipation for motility is in good agreement with that predicted using resistive force theory, indicating that any thermodynamic inefficiencies upstream of the flagellar motor, such as metabolic inefficiencies, are small. We further show that the multiflagellated bacterium *Escherichia coli* dissipates 0.70 µW/mL/OD when in motility buffer, comparable to marine bacteria despite the much lower swimming speed, and hence indicating lower efficiency of swimming. Finally, we show that the energy spent by *Vibrio coralliilyticus* on motility can have major consequences on their survival under starvation, causing a 50 % reduction in survival of a motile wild type relative to a non-motile mutant after 3 weeks of starvation. These results reveal the large costs involved in an individual trait that shapes microbial life in heterogeneous environments.

**Significance:** Many bacteria live in heterogeneous environments where they use motility and chemotaxis to locate nutrient hotspots or hosts amid nutrient-poor waters. The energetics of this search process has not been measured directly. Here we use calorimetric measurements of cultures of swimming and non-swimming bacterial populations to measure the energy expenditure of motility and compare with theoretical predictions. We show that bacterial motility is among the most expensive cellular traits under starvation conditions, accounting for 35% of all metabolic costs. Quantifying the energetic cost of cellular traits helps understanding behavioral trade-offs and energy allocation in natural environments.

---

As famously argued by the physicist Erwin Schrödinger, living systems continuously dissipate free energy in order to overcome the impeding disorder imposed by the second law of thermodynamics (1). All living systems are fundamentally limited by energy and subject to evolutionary pressure to use free energy as efficiently as possible. Therefore, bioenergetics has emerged as a key candidate principle to understand the function, design and evolution of cellular systems and the evolution of cellular traits (2). This requires proper quantification of the energetic resources involved in cellular traits.

Cellular traits have production and operation costs. Advances in genomics, proteomics and metabolomics have enabled more accurate accounting of cellular economics, leading to better understanding of the production costs of cellular traits (3). This in turn has revealed resource allocation trade-offs in bacterial growth and survival (4, 5). The post-translational cost, or the operation cost of cellular traits, can be a major burden to cells but is much harder to quantify. There have been strong conceptual advances in understanding cellular dissipation from the application of non-equilibrium thermodynamics to living systems (6, 7), but these advances are limited by the lack of direct measurements of energetic fluxes, leaving non-equilibrium models restricted to limited cases (e.g., kinetic proofreading, chemotactic receptor sensing) where the processes are characterized in full molecular detail (6, 8, 9). Therefore, new approaches that directly measure the bioenergetic cost of traits will benefit our understanding of the bioenergetics of cellular traits.

Motility is a trait employed by many bacteria to navigate heterogeneous environments. It increases their encounters with isolated hotspots, thereby affecting biogeochemical transformation rates (10, 11), and enhancing encounters with symbiotic partners (12) or host cells, affecting host health & disease (13). The bioenergetic cost of motility has been investigated for a long time in bacteria and archaea (14–20). Despite this attention, quantitative estimates of bioenergetic costs of bacterial flagellar motility vary considerably: from less than 1% (14, 15) of cellular metabolism to about 10% (18, 21). We therefore sought to obtain more direct measurements of the bioenergetic cost of bacterial motility.

Isothermal microcalorimetry is a classic technique that has been extensively used in biochemistry (23), for example to provide most of the reaction enthalpies in the tables of the Handbook of Chemistry and Physics. Recently, there have been pioneering applications of calorimetry to microbiology (24, 25) to quantify heat fluxes during cell growth (26) and starvation (27), but it has never been used to quantify the energetics of an individual bacterial trait. Here, we use calorimetry to quantify the heat flux, and thereby the power dissipation, due to bacterial flagellar motility.

Variation in the estimates for bioenergetic costs of motility can be partly explained by how translational and post-translational costs are weighted. The main focus when it comes to the energetics of bacterial motility has been on its proteomic (or translational) costs, which amount in *E. coli* to a few percent of the proteome. Hence, in well-mixed conditions, cells without motility and chemotaxis genes have a growth rate that is a few percent higher than the wild type (16, 17, 28). The post-translational cost of motility is in the energy that the cell needs to invest to overcome the drag acting upon its body and flagellum as it swims, effectively the cost of operating (rotating) its flagellum or flagella. Because the translation cost is large (about 1000 ATP equivalents per protein molecule (3)) and the motility machinery needs to be produced once per cell cycle, the translational power expenditure depends on the cell cycle duration. During fast growth, the translational cost of motility is dominant over the post-translational cost (10). However, in the natural environment bacteria are often in conditions of starvation rather than active growth (29, 30), as many environments are energy-limited (31, 32). In the absence of growth, translation and transcription of cellular machinery mostly halts, implying that post-translational costs of motility become dominant over translational costs. To quantify the post-translational costs of bacterial motility, we performed calorimetry measurements in the absence of growth comparing motile bacteria with non-motile mutants, and found that in marine bacteria motility requires more than one third of the cell’s entire metabolic budget.

## Results

### Microcalorimetry measurements of energy dissipation due to motility

To measure the power dissipated by motility using microcalorimetry, we used motile and non-motile populations of bacteria of *Vibrio coralliilyticus* YB1, a coral pathogen with a single sheathed polar flagellum. *Vibrio coralliilyticus* has shown persistent motility during growth arrest induced by carbon starvation (22). We constructed a non-motile mutant by deletion of the *motX* gene (Methods), which yields a strain that builds flagella but cannot swim due to the prevention of flagellar motor stator recruitment (33). The stators are responsible for the torque generation required to rotate the flagellum by exploiting an ion gradient (sodium in marine bacteria) (34): failed stator recruitment stops the ion flow and flagellar rotation, thus eliminating the power dissipation associated with motility. We confirmed by microscopy that the mutant was completely non-motile (Supplementary Videos 1 and 2).

To compare the dissipation of the motile and non-motile bacterial populations, we grew both strains in rich medium to early stationary phase and induced carbon starvation by washing the cell suspension in artificial seawater (ASW) without added nutrients, creating non-growing bacterial suspensions in ASW at a concentration of ~ 10^8^ *−*10^9^ cells/mL (OD= ~0.1-1.0). We placed these samples in a microcalorimeter for 6 days (144 h) and measured the heat outflux Φ(*t*) (Fig. 1A). Φ(*t*) typically decreased rapidly over the first ~24 h, until it approximately stabilized in the µW/mL/OD range. We interpret this initial decay as a physiological adaptation phase during which bacteria adjust their metabolism to the starvation conditions in the buffer (Fig. 1B, gray area), as has been observed in other starvation experiments (35). After this adaptation phase, the heat flux of the non-motile mutant Φ(*t*)_ΔMotX_ lay consistently below the heat flux of the wildtype Φ(*t*)_WT_, indicating that motility measurably increased the power dissipation (Fig. 1B). The time-averaged power dissipation of the population due to motility, *P* = ⟨Φ(*t*)_WT_ *−* Φ(*t*)_ΔMotX_ ⟩, was 0.91 µW/mL/OD (Fig. 1C). When normalised to the total energy dissipation of the wildtype, as *P* ^*∗*^(*t*) = (Φ(*t*)_WT_ *−* Φ(*t*)_ΔMotX_)*/*Φ(*t*)_WT_ (Fig. 1D), the power dissipation due to motility plateaued at 39% (*t >* 50 h) and was on average 35% (24 *< t <* 144). This result shows that a substantial fraction of the bacteria’s energy resources during starvation are allocated to motility and that bacterial motility is therefore a costly trait.

**Fig. 1.**
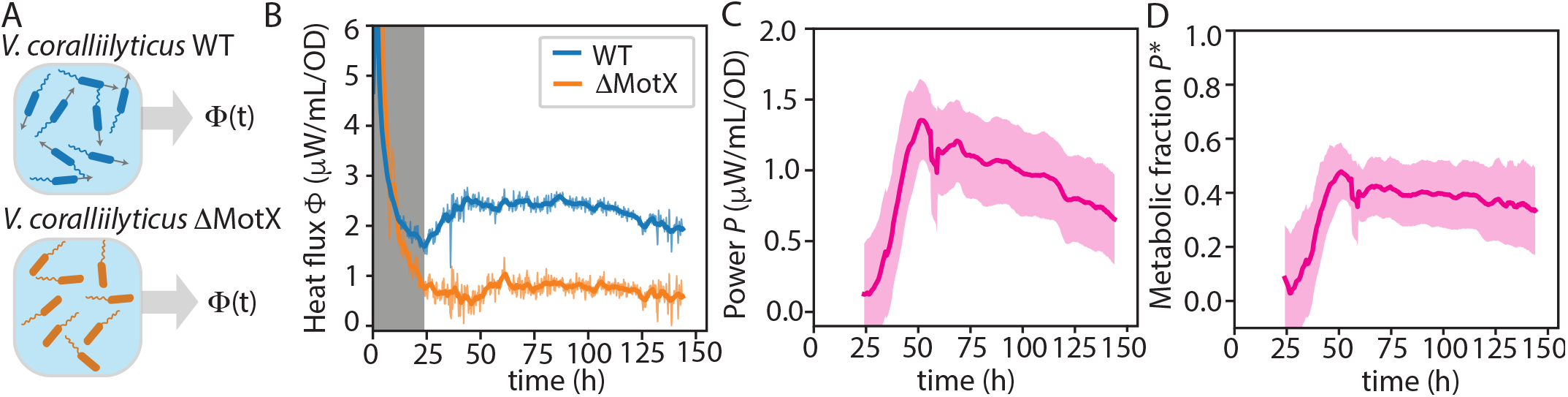
Microcalorimetry measurements reveal large energy expenditure of bacterial motility during starvation. (A) Schematic of a microcalorimetric measurement where the heat flux is measured from a vial with a suspension (OD 0.1–1.0) of either motile (top) or non-motile (bottom) bacteria. (B) Representative time series of heat flux Φ(t) from populations of a marine bacterium *Vibrio coralliilyticus* YB1 (WT, blue) that is motile in the absence of external nutrients (22) and from a non-motile mutant in the same background strain (ΔMotX, orange). A 3 h moving average (colored lines; from two technical replicates) is superimposed over the raw measurements (gray lines). The gray area indicates the physiological adaptation phase to the starvation conditions. (C) The power spent on motility *P* (t), defined as *P* (*t*) = Φ(*t*)_WT_ *−* Φ(*t*)_ΔMotX_, representing the difference in energy dissipation by the motile and non-motile populations. The line and shaded area represent the mean and standard error of the mean from four biological replicates. (D) Motility dissipation as a fraction of total dissipation during starvation, defined as *P*^*∗*^(*t*) = (Φ(*t*)_WT_ *−* Φ(*t*)_ΔMotX_)*/*Φ(*t*)_WT_.

To further support that the difference in power dissipation between WT and the motility-defective *motX* mutant populations was due to cell swimming, we performed an experiment in ASW in which the sodium ions were exchanged 1:1 for potassium ions (Methods). As the flagellar motor of *Vibrio coralliilyticus* is driven by a sodium gradient, exchanging sodium for potassium impairs motility while maintaining the same ionic strength: partial replacement reduces the swimming speed (36). This method does not have any apparent physiological side effects (36). We again measured the difference in energy dissipation between wildtype and mutant using microcalorimetry, for three different sodium concentrations (2, 240 and 479 mM) (Fig. 2A). To determine motility levels, we sampled from cultures grown under identical conditions as in the calorimetry experiments and quantified swimming velocities using video microscopy and particle tracking (Methods; Fig. 2B). We found that the swimming speeds were respectively 7 ±1, 11 ±2 and 25± 16 µm/s (mean ±one std). The calorimetry measurements revealed that the power dissipation due to motility increased with the average velocity of the population, from 0.38 µW/mL/OD in the low-sodium (low-motility) condition to 1.84 µW/mL/OD in the high-sodium (high-motility) condition (Fig. 2B). A linear fit between the energy dissipation and measured mean squared velocity ⟨*P*⟩ = Ω ⟨*v*^2^⟩ yielded a slope Ω = 1.4 *±* 0.4 Pa *·* s *·* µm. These findings show that increasing the swimming velocity of the cells increases the energy dissipation (Fig. 2B).

**Fig. 2.**
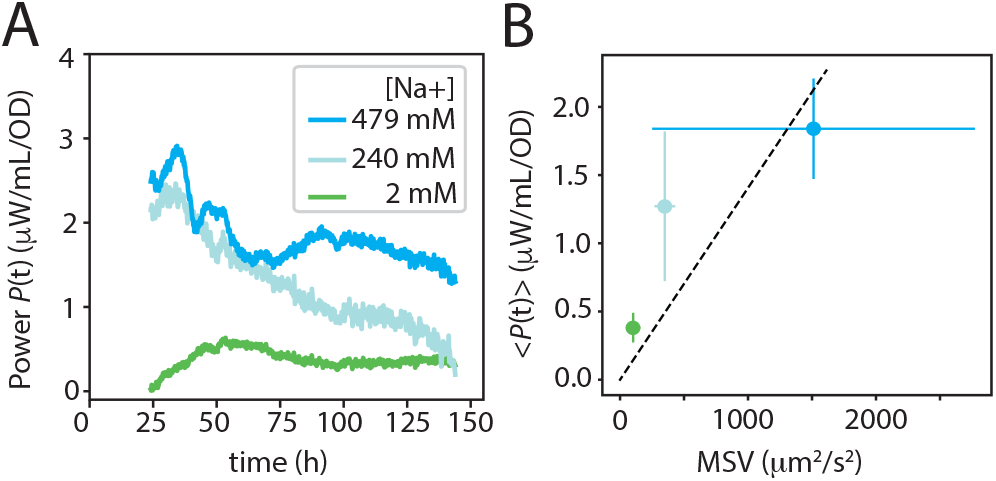
Increasing bacterial swimming velocity increases power dissipation. (A) Calorimetry measurements of motility power for bacterial populations in seawater medium where swimming velocity was manipulated by exchanging sodium, required for flagellar rotation, with potassium (colored lines). Each curve shows the mean of 3 biological replicates. (B) The mean power dissipation *< P >* as a function of the mean-squared velocity (MSV) ⟨*v*^2^⟩, as obtained by video microscopy and particle tracking and averaged over the 24-144 h time interval. Points and error bars represent respectively the mean and standard deviation of three biological replicates. The dashed line represents a linear fit of form *⟨P⟩* = Ω ⟨*v*^2^⟩ with slope Ω = 1.4 *±* 0.4 Pa *·* s *·* µm.

### Inefficiencies upstream of the flagellar filament are small in marine bacteria

We endeavoured to determine the efficiency of the power dissipation due to motility. We determined that a considerable fraction of the total metabolism *P*_*c*_ is associated with motility *P*_*m*_ (Fig. 1 and Fig. 3A). But how efficient is the conversion into propulsive power *P*_*t*_? Three conversions can be identified following the energy fluxes from metabolism downstream to propulsion (Fig. 3A): 1. any loss upstream of the flagellar motor, such as metabolic inefficiencies; 2. loss at the flagellar motor itself; and 3. any loss by rotation of the flagellum (and the counterrotation by the bacterial cell body). These conversions can be described by respective efficiencies of *η*_*m*_ = *P*_*r*_*/P*_*m*_, *η*_*r*_ = *P*_*f*_ */P*_*r*_ and *η*_*f*_ = *P*_*t*_*/P*_*f*_ (Fig. 3A). Our aim is to characterise these efficiencies, to better understand the metabolic costs of motility.

**Fig. 3.**
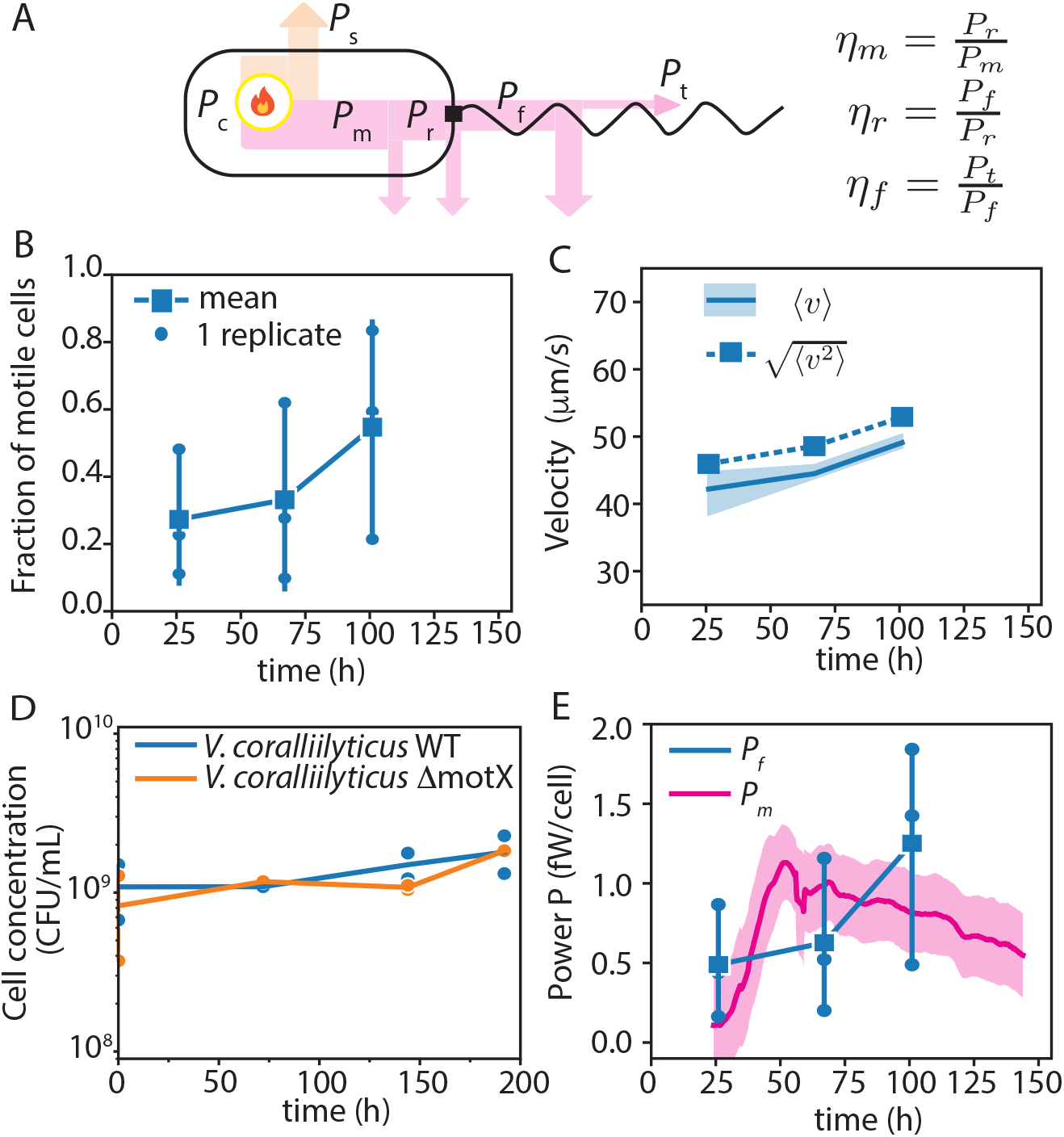
Inefficiencies upstream of the flagellar motor in *Vibrio coralliilyticus* are low. (A) Schematic illustration of the energy fluxes and efficiencies due to bacterial motility. The power generated by metabolism of the cell *P*_*c*_ fuels all cellular processes, including all non-motile and motile functions (*P*_*s*_ and *P*_*m*_, respectively). A fraction of the power dissipated by motility, is channelled to the rotor *P*_*r*_, the bacterial flagellar motor. A fraction of the motor power is transmitted to the flagellum *P*_*f*_, and in turn a fraction of that is converted into thrust *P*_*t*_. At each conversion, a fraction of the power is lost directly due to heat (arrows), and calorimetry measures the total outward heat flux (e.g. the sum of all arrows), equal to *P*_*c*_. (B) Fraction of motile cells quantified from tracking. Squares and error bars represent the mean and standard deviation of three biological replicates (circles). (C) Average velocity (solid line, with shaded error bar) and root-mean-squared velocity (dashed line) of the cells, from the same experiments as panel B. (D) Cell concentration measured using colony counting for wild-type *Vibrio coralliilyticus* (WT, blue) and motility-defective *Vibrio coralliilyticus* (ΔMotX, orange) as a function of starvation time. Each data point represents a single plate (technical replicate). The starting ODs were respectively 1.01 and 1.00. The average cell count for WT was 1.2*·*10^9^ cells/mL/OD. A similar experiment with cell counts measured by flow cytometry yielded 1.7*·*10^9^ cells/mL/OD. (E) Comparison of measured motility dissipation *P*_*m*_ with theoretically estimated motility power requirement *P*_*f*_. Dissipation per cell (purple) was computed by dividing the measured dissipation (Fig. 1C) by the average number of cells (panel D). Motility power requirement (blue) was computed as *P* = (*A*_0_*/η*_*f*_)*f < v*^2^ *>*, where *A*_0_ is the hydrodynamic resistance of the bacterium with its flagellum, *η*_*f*_ the efficiency of the flagellar filament (0.8 %, see main text), *f* the fraction of motile cells (panel B) and *< v*^2^ *>* the average squared velocity (panel C). Squares and error bars represent the mean and standard deviation of three biological replicates (circles).

Previous experiments investigating the energetics of motility have determined the power dissipation of the flagellar filament *P*_*f*_, based on the measured torque exerted on the filament by the flagellar motor (37, 38), and computing its corresponding efficiency *η*_*f*_. However, this does not address any potential losses upstream of the flagellum (*η*_*m*_ *<* 1 or *η*_*r*_ *<* 1). Torque and ion flow measurements indicated that the proton-driven flagellar motor of *E. coli* operates near unit efficiency (*η*_*r*_ *≈* 1) at low rotational frequencies (~ 100 Hz) (39), whereas torque measurements on *Vibrio alginolyticus* indicated that the sodium-driven bacterial flagellar motor (with much higher frequencies, ~ 1000 Hz), operates far from unity (*η*_*r*_ *<<* 1) (40). Inefficiencies upstream of the motor, arising from any loss in metabolism, for example from restoring the transmembrane ionic gradient that fuels the flagellar motor, have not been considered for flagellar motility, but have been considered in other systems. For example, in the case of the energy release of a methyl group in bacterial chemotaxis, an efficiency of 20% has been used to compute the required power dissipation (41). Similar efficiencies in the conversion of chemical energy to the ion gradient that powers bacterial motility would imply a much higher power dissipation due to motility than measurements of *P*_*f*_ suggest. Using calorimetry, theoretical predictions from resistive force theory and swimming parameters, we can provide bounds on the efficiencies *η*_*r*_ and *η*_*m*_.

To quantify the efficiencies upstream of the flagellar motor, we need to compare total power dissipation (due to motility) per cell *P*_*m*_ with the power dissipation at the level of the flagellar motor, *P*_*f*_. The efficiency upstream of the flagellar motor equals *η*_*u*_ = *η*_*m*_*η*_*r*_ = *P*_*f*_ */P*_*m*_. To estimate *P*_*f*_, we use resistive force theory. The population-averaged motility power requirement can be computed as

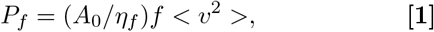

where *A*_0_ is the resistance coefficient of the bacterium, *η*_*f*_ the efficiency of the flagellar filament, *f* the motile fraction of cells and *< v*^2^ *>* the mean squared velocity (MSV) (42). To obtain the swimming parameters, we performed experiments in vials that mimic the conditions in the calorimeter and sampled to perform video microscopy and cell tracking: the motile fraction (defined as the time-averaged fraction of cells with an average swimming speed larger than 12 µm/s, Ref. (22)) was 0.38 ± 0.19 (mean ± std from three biological replicates; Fig. 3B), the mean swimming speed of motile cells was 45±1 µm/s, and the root mean squared velocity (RMSV) was 49 ± 1 µm/s (Fig. 3C).

The resistive force constant *A*_0_ can be computed from the cellular and flagellar morphologies (38). We extracted morphological parameters of the same strain used in experiments after 24 h of carbon starvation, based on previously acquired scanning electron microscopy data (22). We found a cell length and width of 1.49 ± 0.10 µm and 0.43 ± 0.01 µm (mean ± s.e.m.), respectively, and a flagellar length, amplitude, and pitch of 4.1 ± 0.14 µm, 0.46 ± 0.03 µm, and 1.80 ±0.05 µm, respectively (full morphological parameters for 13 cells are in Supplementary Table 1). We computed *A*_0_ by approximating the cell head as a weakly-elongated spheroid, with the measured cell length and width used for the long and short axis, respectively, yielding *A*_0_ = 6.56 ± 0.23 mPa· s ·µm.

For the flagellar efficiency *η*_*f*_, there is a body of work that expresses the efficiency in terms of the flagellar and cellular morphologies using resistive force theory (RFT) (Methods). Using the RFT formalism of Gray and Hancock (43) and our measured morphological parameters, we find an efficiency *η*_*f*_ = 0.80%. This efficiency is within the range of previous estimates for *E. coli*, which range from 0.2% (38) to 3% (42).

To obtain a per-cell estimate of the power dissipation due to motility we determined *P*_*m*_ =*< P > /N*_*c*_, where *< P >* is the power per volume per unit biomass (OD_600_) measured by calorimetry averaged over the 24-144 h time window (Fig. 1C) and *N*_*c*_ the cell concentration per unit biomass. To obtain an estimate for *N*_*c*_ in the calorimetry experiment, we sampled cells from starved cultures to estimate cell counts (from plating) and we found that both the motile and non-motile populations had an approximately constant cell concentration of *N*_*c*_ =1.2 ×10^9^ cells/mL/OD over the time period (based on an initial density of OD_600_ = 1.0). We used *N*_*c*_ to compute the power dissipation per cell due to motility (Fig. 3E, purple curve), which yielded *P*_*m*_ = 0.76 ± 0.28 fW (1 fW = 10^*−*15^ W). Using Eq. 1, we thus estimate a power dissipated by the flagellar motor of *P*_*f*_ = 0.79 ± 0.49 fW. Hence the efficiency *η*_*u*_ = *P*_*f*_ */P*_*m*_ ≈1, and this constrains the efficiencies *η*_*m*_ and *η*_*r*_ to be close to unity as well. Together, this analysis shows that the motile *Vibrio coralliilyticus* loses energy through the rotation of the flagellar filament, consistent with what was described previously (38, 42), but any losses upstream of the flagellum, for example due to metabolic losses, are minor. This means that the bacterial flagellar motor of *Vibrio coralliilyticus* operates with near-unit efficiency, unlike what might have been expected based on torque measurements on *Vibrio alginolyticus* (40). It also suggests that any potential losses due to internal friction between flagellar sheath and filament (44, 45) are minor.

Apart from RFT in the original Gray and Hancock formalism (43), there are alternative descriptions of the hydrodynamics of flagellar rotation, notably the RFT formalism of Lighthill (46), and slender body theory (SBT) (47). Using either Lighthill-RFT or SBT, we find values of the efficiency of respectively 0.25% and 0.40%. These flagellar efficiencies would predict an upstream efficiency *η*_*u*_ that exceeds unity, of respectively 2.9 and 2.1 (Fig. S4). Such unphysical efficiencies suggest that these theories overestimate the power dissipated by the flagellar motor in this system (SI Appendix, Fig. S4 and S5, Supplementary Text).

### Power dissipation due to motility in *E. coli*

We then explored the generality of the motility expenditure by performing experiments with *Escherichia coli. E. coli* typically uses 3–4 flagella to swim with a speed of approximately 25 μm/s (48–50), much lower than typical marine bacteria (51). In the absence of an energy source, starving *E. coli* gradually lose motility within one day (22) and slow down chemotactic signal processing (41), making it difficult to study motility in the absence of growth. However, motility in the widely used strain RP437 can be investigated by exploiting its auxotrophic limitations of 4 amino acids: this strain remains motile in motility buffer lacking these amino acids, where the cells do not grow but an energy source is provided (52, 53). We thus performed a microcalorimetry experiment in motility buffer, comparing RP437 with two flagellar stator mutants, (ΔMotA and ΔMotB), which are non-motile. Stator-associated mutations are often employed to eliminate flagellar motion without disruptive effects to physiology (54) and prevent any physiological changes due to flagellar-dependent gene expression (55).

We measured a power dissipation due to motility of *P*_*EC*_ = 0.70μW/mL/OD when comparing RP437 to the ΔMotA mutant (Fig. 4A) for the same time window (24-144 h) as used for marine bacteria and 0.78 μW/mL/OD when averaged over the entire experiment except the initial 24-h acclimation period (24 *< t <* 240 h). Comparing the power dissipation of RP437 to the other non-motile mutant (ΔMotB) yielded a similar value of 0.73 μW/mL/OD (Supplementary Fig. S3). This amounted to 25–40 % of cellular metabolism 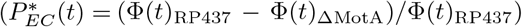of *E. coli* in the motility medium, with the value slowly increasing over time through that range (Fig. 4B). This shows that despite the differences in metabolism, medium conditions and flagella architecture, *E. coli* and *V. coralliilyticus* dissipate a similar fraction of their metabolism to motility.

**Fig. 4.**
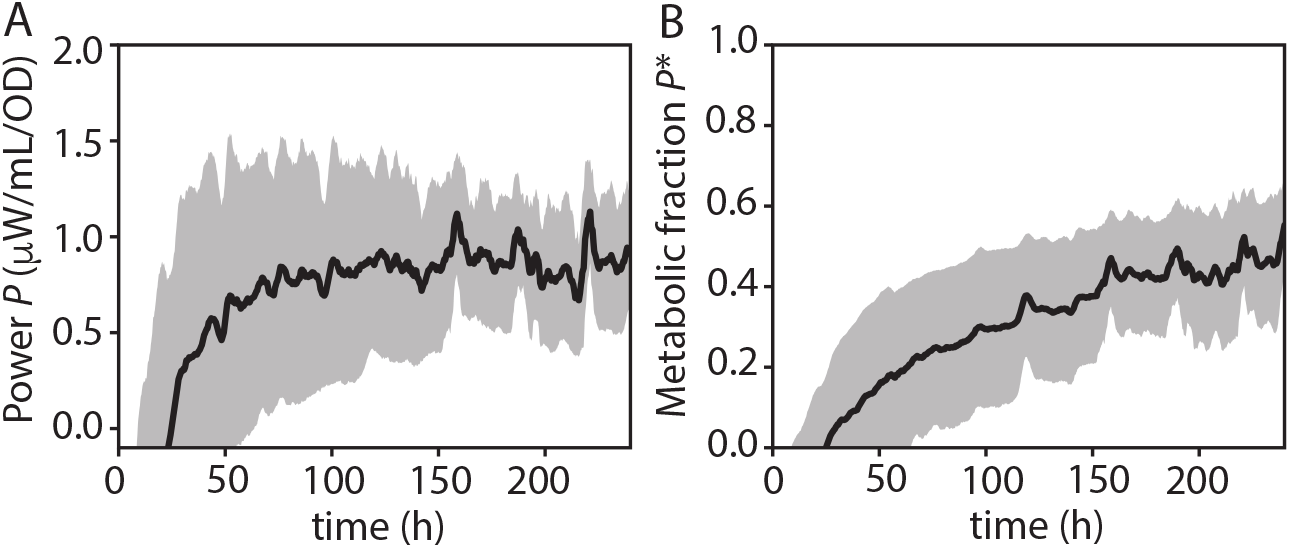
Energetics of motility in *E. coli*. (A) Motility power dissipation in *E. coli* RP437, determined as the difference in measured heat flux between motile strain RP437 (WT for chemotaxis) and non-motile strain RP6647 (ΔMotA), in motility buffer. Due to the auxotrophic limitations of RP437 and RP6647, these strains do not grow in motility buffer, but RP437 continues swimming. The average of three biological replicates with 3 h moving average (black line) is shown together with the standard error of the mean (gray shading). (B) Fraction of dissipation due to motility in *E. coli*, defined as the motility power dissipation (panel A) divided by the total dissipation of the wild-type strain (RP437). Curve and shading as in A. The corresponding curve for *Vibrio coralliilyticus* is shown in Fig. 1D.

### Dissipation due to motility in homogeneous environments creates a survival deficit

A cell’s energy utilization during starvation is particularly important because a lack of energy may expedite cell death (56). We did not observe pervasive cell death in the first week of starvation of *Vibrio coralliilyticus*, as cell numbers remained approximately constant (Fig. 3D). To quantify the impact of the power dissipation due to motility on bacterial survival during starvation over longer timescales, we performed an experiment in which *V. coralliilyticus* cells were starved for up to 4 weeks at room temperature. As potential cell lysis after cell death releases nutrients, we kept the cultures under continuous shaking to homogenise the environment and exclude any potential benefit from motility and chemotaxis during the starvation period. The cultures were sampled regularly for colony counting. We then quantified bacterial survival as the count of colony forming units (CFU) for both WT and motility-defective (ΔMotX) populations (Fig. S1). The relative survival was quantified as CFU_WT_*/*CFU_MotX_, where CFU_WT_*/*CFU_MotX_ *<* 1 implies an excess death rate associated with active motility during starvation. We observed that the ratio CFU_WT_*/*CFU_MotX_ was close to unity during the first 10 days of starvation (Fig. 5), but after two weeks of starvation the value began to decrease, reaching a value of CFU_WT_*/*CFU_MotX_ = 0.5 after three weeks of starvation. This shows that the energetic cost of motility results in a substantial survival deficit during starvation in well-mixed environments (i.e. where there is no benefit), albeit with a two-week delay.

**Fig. 5.**
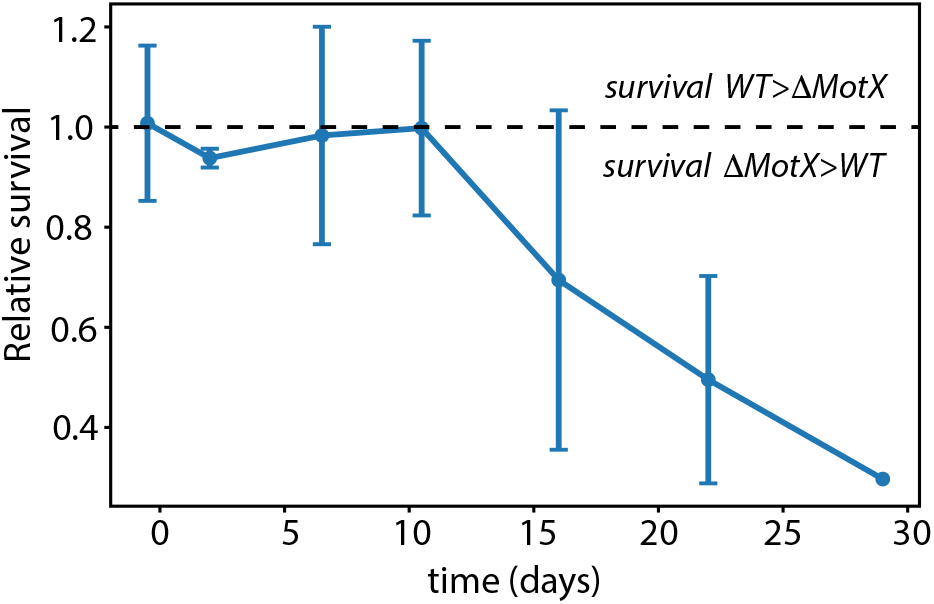
Energy expenditure on bacterial motility in well-mixed environments results in a survival deficit during starvation. Points show the relative survival of motile cells as a function of time, quantified as the ratio of CFU counts in samples of WT (motile) and ΔMotX (non-motile cells) at each time point. The horizontal dashed line indicates equal survival probabilities of motile and non-motile cells. The blue line and error bars represent the mean and standard deviation of three biological replicates.

## Discussion

We have here provided the first direct measurements of the energetic cost of a bacterial trait and its comparison to the native metabolism of the same population. For marine bacterium *Vibrio coralliilyticus*, we find that on average motility carries an energetic cost of 0.76 fW/cell. However, this estimate is based on a population where not every cell swims. Given an average motile fraction of 0.38, an estimate based on only motile cells would yield 2.0 fW per motile cell, equivalent to a turnover of 2.5 *×* 10^4^ ATP molecules per second. The majority of this expenditure is due to the inefficiency of converting flagellar motor rotation into propulsion, but our calorimetry results suggest that other potential inefficiencies in *Vibrio coralliilyticus* are low.

The dissipation due to motility is considerable relative to the metabolic activity during bacterial growth arrest (Fig. 1D). For comparison, during fast growth *Vibrio coralliilyticus* WT dissipates 1–10 mW/mL/OD (Fig. S2), a thousand-fold greater value compared to the metabolism during starvation. Therefore, compared to growth — for which the majority of the costs are associated with protein synthesis — flagellar rotation reflects only a small fraction of metabolism. During growth, a more appropriate metric for the demand for cellular resources is the fraction of its proteome devoted to motility – typically equal to a few percent of cellular proteome, and therefore total metabolism (17, 57) – and not the metabolic demand of flagellar rotation. However, given the expenditure on motility of 35% due to flagellar rotation alone, motility is likely the most expensive trait during starvation.

Our measure of basal energy dissipation rate, or maintenance energy, of ~ 1 fW/cell (or 1 · 10^4^ ATP/cell/s), in both growth-arrested *Vibrio coralliilyticus* as well as *E. coli*, is comparable to previous estimates based on the minimum carbon source necessary to postpone cell death during carbon starvation in *E. coli* (56). However, this is still nearly ten-fold higher than the minimum measured metabolism level of ~ ·10^3^ ATP/cell/s reported to date (35), which is in turn close to the theoretical minimum dissipation required for ion homeostasis based on intrinsic membrane permeability (58), the main challenge for cellular survival (59). Hence, there seems to be some behavioral freedom in the baseline metabolism of bacteria during starvation and calorimetry could potentially be used to quantify those differences.

We estimate that the power dissipation due to motility represents 35 % of metabolism during starvation, but this value could be much higher or lower depending on motility parameters. There are reports of bacteria in seawater that swim much faster than the ~ ·50 µm/s that we report here (60). If we assume the same baseline metabolism and scaling as in eq. 1, a bacterium (temporarily) swimming with a speed of 200 µm/s would spend 97.5 % of its power on motility (or hypothetically 99.8% with a maintenance energy close to the theoretical minimum of 1000 ATP/s). Such motility speeds are likely not sustainable for bacteria, and indeed measured speeds across marine bacteria during starvation are lower than the 50 µm/s measured for *Vibrio coralliilyticus*(22).

It is interesting that the power dissipation due to motility in *E. coli* is 77 % of the dissipation measured for *Vibrio coralliilyticus*, corresponding to a 1.3-fold decrease. Given that *Vibrio coralliilyticus* swims nearly twice as fast as *E. coli*, a four-fold decrease in motility energy dissipation would be expected (if cellular morphology were the same), given the dependency of *P*_*m*_ on *v* in Eq. 1. The fraction of motile cells *f* is typically high in *E. coli* populations, approximately 0.8–0.9, (Ref. (49, 57), compared to 0.38 in *Vibrio coralliilyticus*), but even a motile fraction of 1.0 in *E. coli* would not fully explain this difference. Given the similarity in *P*_*m*_, this means that one or more of the downstream efficiencies (*η*_*m*_, *η*_*r*_ or *η*_*f*_) are lower in *E. coli* compared to *Vibrio coralliilyticus*. A possible explanation is that in multi-flagellated bacteria there are additional dissipation losses, for example due to inter-flagellar resistance (61) or due to bacteria operating their flagellar motor in the low-torque, high frequency regime where the flagellar motor is less efficient (62). This is consistent with single-flagellated marine bacteria having a flagellar arrangement that is optimized for speed, with a lower energy expenditure per unit velocity compared to multi-flagellated enteric bacteria. The motility of multi-flagellated bacteria could be optimized for something other than speed, such as torque generation in highly viscous media (62) or navigation in porous environments (63).

In this work, we have quantified the cost of bacterial motility. In natural settings, these costs are expected to be offset by the benefits to the motile cell. Bacterial motility is important in patchy environments, where nutrients are distributed heterogeneously (64). One example could be the colonization of marine particles, where successful colonization creates many offspring, and even if only a small fraction of searching bacteria successfully encounter a particle, this may offset the cost for the population (65). Another area where bacteria may benefit from motility in the absence of growth is the search for hosts by pathogens. Motility has been shown to be important for *Vibrio coralliilyticus* in its infection of coral hosts (66). This implies that inhibiting bacterial motility around corals or other marine hosts may represent an approach for pathogen protection. The energetics of motility for human or animal pathogens could be a fertile direction for future research.

We have shown previously that *Vibrio coralliilyticus* fuels its motility by consuming internal energy reserves (22). The consumption of these energy reserves does not have an immediate effect on survival. However, long-term survival is impacted because the energy reserves required to survive are already spent on motility. Nonetheless, a relative survival deficit of about 0.5 of motile cells compared to motility-defective cells after 3 weeks seems minor. Considering again a foraging scenario similar to particle foraging, if the enhanced exploration yields on average a single additional division over 3 weeks, this would offset the motility cost in terms of the decrease in survival. Given that motility boosts bacterial encounters with nutrient hotspots by 100–1000 fold (67, 68), this implies that in many scenarios motility is worth the energetic investment. But since many bacterial species in fact do not swim when starved (22), this suggests that while bioenergetics is important, the cellular decision to be motile or not may also include non-energetic factors.

## Materials and Methods

### Bacterial strains and genetics

Experiments with marine bacteria were performed with *Vibrio coralliilyticus* YB1 (ATCC BAA-450, Ref. 69). *E. coli* experiments were performed on RP437 (*thr(Am)-1 leuB6 eda-50*, a derivative of K-12 and considered wild type for chemotaxis (70)), and non-motile mutant strains RP6647 (*motB*Δm14-1 (lac)Δu169, Ref. 71) and RP666 (*motA*Δm12-14 (lac)Δu169, Ref. 71). All *E. coli* strains were kind gifts of John S. Parkinson, University of Utah.

A motility-deficient mutant of *Vibrio coralliilyticus* YB1 was created by deleting the *motX* gene (which encodes the motor protein MotX (NCBI EEX30621)), by homologous recombination using a mutagenesis plasmid. To create the plasmid, the primers B450-motX-up-EcoRI-F (ATATATGAATTCtcatatcggttacatccttggtatca, all primer sequences read 5’ to 3’) and B50-motX-up-OEX-R (ACCAGAAAGTATCGCGTAACTTTCT-TAGCTTCATtgtctgcactcttt) were used to amplify approximately 800 bp of DNA upstream of *motX* via PCR, and the primers B450-motX-down-OEX-F (GAAGCTAAGAAAGTTACGCGATACTTTCTGGTAAagcgataatctgac) and B450-motX-down-XbaI-R (ATATATACTAGTaatacgactgaaactctgcagaagatt) were used to amplify approximately 800 bp downstream of *motX*. The two products were purified and used as products for overlap extension PCR (72) together with primers B450-motX-up-EcoRI-F and B450-motX-down-XbaI-R. The approximately 16 kb product was purified and then digested with the restriction enzymes EcoRI and XbaI. The product was then cloned into the suicide plasmid pSW4426T (73) digested with the same restriction enzymes to create the plasmid pBU260. To create the knockout deletion, a previously described protocol using homologous recombination was used (66). Briefly, the mutagenesis plasmid pBU260 in *E. coli* β 3914 (73) was introduced into *V. coralliilyticus* YB1 via conjugation. After two successful rounds of recombination, potential mutants were initially screened for motility using soft-agar chemotaxis assays (66) followed by PCR using the primers B450-motX-up-EcoRI-F and B450-motX-down-XbaI-R. All confirmed motX mutants were screened for any growth defects by comparing growth over time against the parental strain (YB1).

### Cell culture and starvation protocol

For experiments with Vibrio, strains were grown in 2216 Marine Broth from a glycerol stock stored at −70 C. Cells were cultured for 24 h and then washed three times by centrifugation (4500 *× g*, 5 min) in artificial seawater (Instant Ocean, ‘ASW’) or for some experiments in defined seawater in which the concentration of NaCl was changed by exchanging sodium with potassium (Experiments in Fig. 2, Supplementary Table 2). No nutrients or buffers were added to the seawater, thereby inducing nutrient starvation in the bacteria. The optical density was measured using a photospectrometer (brand, place) and adjusted to OD by further dilution in ASW if necessary, minimizing variation in OD between WT and ΔMotX cultures in the same calorimetry experiment.

For the experiments in which swimming speed was manipulated by changing the sodium concentration, the cells were washed one time in Synthetic Ocean Water (SOW), a defined salt mixture from the widely used Aquil medium (74), preparing without H_3_BO_3_, NaF and EDTA. The sodium concentration in the medium was changed by replacing NaCl with KCl, and replacing Na_2_So_4_ with K_2_SO_4_ at equal Na+ molarity, yielding Na+ concentrations of 2 mM (low sodium condition, due to NaHCO_3_), 240 mM (medium sodium) and 479 mM (high sodium).

For experiments with *E. coli*, cells from a glycerol stock stored at −70 C were cultured in Tryptone Broth for 24 h then washed in motility buffer, a minimum motility medium [EDTA, L-methionine, KPO_4_, 10 mM lactic acid, pH 7.0]. Due to the auxotrophic limitations of strain RP437, motility buffer does not support growth (70), but cells do not experience energy limitation due to the presence of lactic acid. In absence of an energy source, *E. coli* can sustain motility for a few hours due to an endogenous energy source (75, 76), but most cells lose motility within approximately 24 h (22).

### Isothermal microcalorimetry

After washing the cultures, the bacterial suspensions were adjusted to ensure that the strains were tested at similar optical densities (OD) at 600 nm in the same experiment (OD 0.4–1.05 for *Vibrio*, 0.15–0.41 for *E. coli*, list of optical densities for each experiment in Table S2). From each culture, 2.9 mL was placed in autoclaved glass calorimetry vials. The vials were immediately closed to be airtight using crimp caps with pre-assembled butyl-rubber septum, leaving approximately 1.1 mL of air available above each culture. The closed vials were then loaded in a TAM48 calorimeter (Waters/TA instruments, Newcastle, DE) and placed in thermal equilibration position for 15 min. After thermal equilibration, the samples were lowered into the measurement position and data acquisition started after 45 min according to the manufacturer’s instructions. To compare power dissipation between experiments, heat fluxes were normalised to both volume and OD. OD was used as a proxy for dry biomass; normalising metabolic rates in this way is standard in quantitative bacterial physiology studies (77).

### Microscopy and cell tracking

Microscopy and tracking was performed as described previously (22, 78). Cells were placed in a chamber made by sealing a rubber spacer between a standard microscopy slide and a coverslip, and imaged at 30 fps and 10X magnification on a Nikon Ti-2 phase contrast microscope. To facilitate tracking, the cells were diluted 1/100 into ASW and imaged within 15 min of being placed in the chamber. Tracking and trajectory analysis was performed using the Trackpy package (79).

### Cell counting

Colony counting was performed by sampling from the starvation culture after serial dilution in ASW, then plating 180 µL culture on 2216 Marine Broth with 1.5% agarose, that was prepared on the same day by autoclaving, plate poring and drying of 30-60 minutes. Plates were sealed with parafilm to prevent drying and cultured at 27 C in an incubator. Plates were photographed and colonies counted manually 20–28 h after inoculation. Plate counts were typically 30– 300 colonies per plate. Plates with less than 10 counts were excluded from the analysis.

For one of the replicates in calorimetry, the cell number at the onset was also measured by flow cytometry. This yielded 1.7*·*10^9^ cells/mL/OD for *Vibrio coralliilyticus* YB1 and 5 *·* 10^8^ cells/mL/OD *E. coli*.

### Resistive Force Theory

Resistive force theory (RFT) was used to estimate the power dissipated by the bacteria as a result of their motility. Assuming no hydrodynamic coupling between cell body and flagella, the forces (**F**) and torques (***τ***) acting on the cell body and flagellum are additive and each described by linear equations of the form

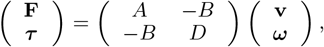

where **v** and ***ω*** indicate the velocity vector and the angular velocity vector, respectively, and *A, B, D* are the resistance coefficients associated with translation (*A*), rotation (*D*) and roto-translational coupling (*B*). We will use a subscript 0 to refer to cell-body quantities, and no subscript for flagellar quantities. We set our reference frame such that, for a left-handed flagellum, all resistance coefficients are positive, *v >* 0, the angular speed of the flagellum is *ω <* 0, and that of the counter-rotating body is *ω*_0_ *>* 0. In bacterial locomotion, power (*P*) is generated by the rotation of the flagellar motor, but only a fraction *ε* (usually small) of this power output is converted into propulsion: *P*_prop_ = *εP*. The propulsive power requirement in the Stokes limit is determined by the drag force acting on the cell body:

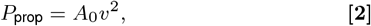

where *A*_0_ is the translational resistance coefficient of the cell body. The propulsive efficiency *ε* is, by the definition, the ratio of the propulsive power requirement (*P*_prop_) and motor power output (*P* = *τ* |*ω − ω*_0_|). At low reynolds number, according to RFT, it can be expressed as a function of only geometrical parameters of the cell body and flagellum (42)

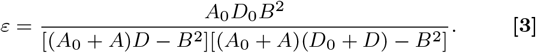

If bacterial morphology and swimming speed are known, it is then possible to estimate the total power output *P* = *A*_0_*v*^2^*/ε* without direct measurements of motor torque.

Flagellar and cell body morphology data for *V. coralliilyticus* YB1 were obtained from SEM images of cells starved for 24 h as reported previously (22). Using these data, we measured cell length *L*, cell width *W*, axial length of flagella Λ, amplitude of flagellar helix *R*, and flagellar wavelength (or pitch) *λ*. We did not measure the flagellar cross-section radius, but assumed it to be *r* = 16 nm as reported previously for *V. alginolyticus* (80) with a similar sheathed polar flagellum. The cell body was described as a slightly eccentric spheroid (*L* ≳ *W*), for which the translational and rotational resistance coefficients along the major axis are (81)

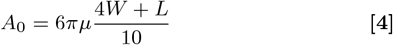

and

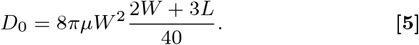

This low-eccentricity approximation is more adequate to describe the YB1 cells in our experiments (with aspect ratios between 1.9 and 5) than the more commonly used slender ellipsoid approximation, which instead only becomes accurate for aspect ratios larger than ~5. In the imaged YB1 samples, we obtained body translational resistance coefficients *A*_0_ ranging between 5.3 and 8.1 µm mPa s with mean value 6.6 µm mPa s, and rotational resistance coefficients *D*_0_ between 0.36 and 1.1 µm^3^ mPa s with mean value 0.69 µm^3^ mPa s.

In RFT, the resistance coefficients for the flagellum are (38)

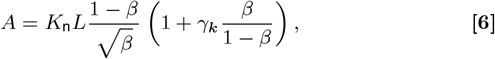

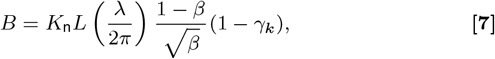

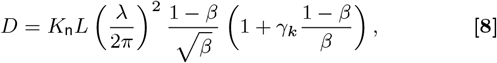

where *β* = (cos Ψ)^2^, with Ψ = arctan(2*πR/λ*) the flagellar pitch angle. *K*_n_ is the perpendicular resistance coefficient, and *γ*_*k*_ = *K*_t_*/K*_n_, where *K*_t_ is the tangential resistance coefficient.

*K*_n_ and *γ*_*k*_ are not fully analytically defined for flagella, since they depend on an empirical parameter for which different approximation schemes have been proposed (see Supplementary Table 1 in Ref. (19) for a summary). Following the nomenclature of Ref. (19), in our study we used the approximation schemes “GH” (the original Gray–Hancock formulation, with 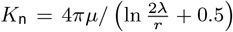 and 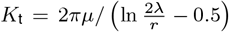 and “LH07” (a modified version of the Lighthill formulation, with 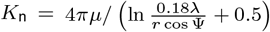 and constant *γ*_*k*_ = 0.7), which provided respectively the upper and lower bound to the estimate of the propulsive efficiency in our geometrical regime.

For both RFT schemes we estimated the corresponding set of resistance coefficients (*A, B, D*) and from these we estimated the propulsive efficiency following Equation 3. Additionally, we calculated these resistance coefficients using Slender Body Theory (see Supplementary Information). We expected *ε* to correlate negatively with cell length *L*, but, interestingly, we observed instead a positive correlation, arising from the covariation of *L* with other geometrical properties of the bacteria (Supplementary Figure 5). For the GH RFT scheme, we estimated *ε* between 0.45% and 1.0% with a mean value 0.80%. For the LH07 scheme, *ε* was nearly 4-fold lower, with values between 0.12% and 0.39% and a mean value of 0.27%. SBT yielded intermediate efficiency values between 0.21% and 0.54% with mean 0.40%.

For the GH RFT scheme, the effective energy dissipation coefficient, Ω = *A*_0_*/ε*, ranged from 0.7 to 1.2 µm Pa s with an average value of 0.8 µm Pa s. For the LH07 scheme, Ω values ranged between 1.8 and 4.4 µm Pa s with an average value of 2.6 µm Pa s. For SBT calculations, we obtained Ω values ranging from 1.3 to 2.5 µm Pa s with average 1.7 µm Pa s.

## SI Datasets

Supply .xlsx, .csv, .txt, .rtf, or .pdf files. This file type will be published in raw format and will not be edited or composed.

## ACKNOWLEDGMENTS

We thank Isobel Short for help in the lab, Sandy Parkinson for sharing strains, Ela Burmeister for logistical assistance, and Russell Naisbit for scientific editing. We also express thanks to Laurens Keegstra for help with colony counting. This work was supported by a Royal Society Newton International Fellowship (NIF *\*R1 *\*253229 to J.M.K.), the Simons Collaboration on Principles of Microbial Ecosystems (PriME; grant 542395FY22 to R.S.), the Swiss National Science Foundation (grant 205321 207488 to R.S.), NCCR Microbiomes, a National Center of Competence in Research funded by the Swiss National Science Foundation (grant numbers 51NF40 180575 and 51NF40 225148 to R.S.) and a Gordon and Betty Moore Foundation Symbiosis in Aquatic Systems Initiative Investigator Award (grant GBMF9197 to R.S.; https://doi.org/10.37807/GBMF9197).

