## Supplementary Information File for "Microcalorimetry reveals bacterial motility as an expensive trait during starvation"

**J.M. Keegstra, O. Braissant, R. Foffi, O. Mueller, V.J. Orphan, and R. Stocker**

**Corresponding Authors: Keegstra and Stocker.**
****

##### This PDF file includes:

Supporting text
Figs. S1 to S5
Tables S1 to S2
Legends for Movies S1 to S2
SI References

##### Other supporting materials for this manuscript include the following:

Movies S1 to S2

### Supporting Information Text

#### 1. Detailed comparison between RFT and SBT

In addition to RFT calculations, we also computed the resistance coefficients of the bacterial flagella using Lighthill's formulation of Slender Body Theory (SBT) (1). Briefly, SBT represents a flagellum as a distribution of Stokeslets and source dipoles arranged along its centerline, yielding an integral equation that describes the force per unit length acting on the surrounding fluid. This approach by construction takes into account long-range hydrodynamic interactions between flows induced by different parts of the flagellum, interactions which are instead neglected by RFT. By evaluating the thrust, torque, and drag on the flagellum, one can then reconstruct the flagellar resistance coefficients, if it is assumed that flagellum and cell body are not hydrodynamically interacting. To perform this computation, we used a Python port of the Matlab code developed by Rodenborn *et al.* to compute the thrust, torque and drag acting on an isolated flagellum (2). The Matlab codebase was translated to Python using Claude Code (version 2.1.119, model Opus 4.7) and mathematical consistency with the original implementation was then manually verified.

Our SBT calculations yielded estimates of propulsive efficiency that are lower than the efficiency estimated by the GH RFT scheme, but higher than the efficiency estimated by the LH07 scheme (Fig. S4, values reported in Main Text – Methods).

RFT is a local approximation of SBT, therefore one would expect SBT to provide a more accurate description of swimming hydrodynamics and therefore of energy consumption (1, 3). For what concerns RFT, none of the different approximation schemes is in general more accurate than the others, and different authors have favored different schemes. Chattopadhyay *et al.* used optical traps to measure the thrust and torque generated by swimming *Escherichia coli*, from which they derived the resistance coefficients and therefore the propulsive efficiency; their estimates of the resistance coefficients were in better agreement with the GH scheme (4). Lisevich *et al.* measured swimming speed and flagellar rotation frequencies of *E. coli* in bulk fluid via high-speed video-microscopy and found that RFT descriptions based on the original GH or LH schemes were systematically overestimating bacterial swimming speed, whereas good quantitative agreement was found with the modified LH07 scheme (5). Johnson has shown with numerical computations that GH RFT tends to consistently underestimate energy expenditure (hence overestimating propulsive efficiency) compared to SBT (3). Spagnolie and Lauga collected geometrical parameters from the literature for all flagellar polymorphs of both peritrichous and polar flagellar configurations and reported that LH RFT can significantly underestimate the efficiency of different flagellar polymorphs and is generally not capable of identifying the most efficient polymorphic form, whereas the non-local interactions embedded in SBT are required to correctly describe propulsive efficiency (6). Rodenborn *et al.* compared the GH and LH RFT schemes against SBT and experiments of macroscopic swimmers in highly viscous fluids, and found that the accuracy of each RFT scheme varies widely across the parameter space, whereas SBT (both in Lighthill's and Johnson's formulations) generally captures the essential physics of swimming with high accuracy (2).

We note that we did not perform a full SBT calculation of the coupled flagellum-body system, but only used SBT to estimate resistance coefficients ($A$, $B$, $D$) for an isolated flagellum and assumed no hydrodynamic interaction between flagellum and cell body. The better agreement of the GH RFT scheme with our experimental measurements then suggests two possibilities:

- The GH approximation scheme overestimates the efficiency by a factor that, in our geometrical regime, is fortuitously similar in magnitude to what would be obtained by fully accounting for the flagellum-body coupling.

- Some of the parameters used in our calculations do not exactly represent the morphology or motile behavior of the cells in the calorimetry experiment, leading to a slight overestimate of power consumption in the SBT calculations; for instance, there may be distortions of flagellar pitch and amplitude due to physical interactions with the substrate mesh in SEM images, or the tracking experiments might be overestimating swimming speed and/or motile fraction due to different environmental conditions from the calorimetry experiments.

However, Chattopadhyay and Wu have highlighted that, while flagellum-body interactions tend to increase propulsive efficiency, they yield a correction that is approximately 2% for bacteria with body sizes comparable to the YB1 used in our experiments (7). Therefore, the neglected flagellum-body interactions cannot explain the nearly 2-fold discrepancy observed between our calorimetry measurements and the SBT estimates. Differences in morphological or swimming parameters between the calorimetry and tracking experiments then appear as the most likely explanation. Another source of error could be due to body shape, which we assumed to be an ellipsoid, whereas Vibrios tend to have a twisted helical-like appearance; similar body shapes, usually found in *Helicobacter* species, have been reported to increase swimming efficiency, although their contribution was limited to an enhancement of at most $10 \sim 20\%$ (8, 9), so the effect of cell morphology on energetics is small (10).

64 ## 2. Supplementary Figures

**Fig. S1. Cell concentration as measured by colony counting** as a function of starvation time in days (d). Colony counting was performed on standard 2216 marine broth plates after serial dilution in ASW (Methods), yielding the concentration of colony forming units (CFU) per mL. Shown are the counts for 3 distinct biological replicates (1: pink, 2: purple, 3: black) of the two genotypes WT (solid) and ΔMotX (dashed). The shaded region and the error bars indicate the 95 CI from technical replicates (multiple plates from the serial dilution), where available.

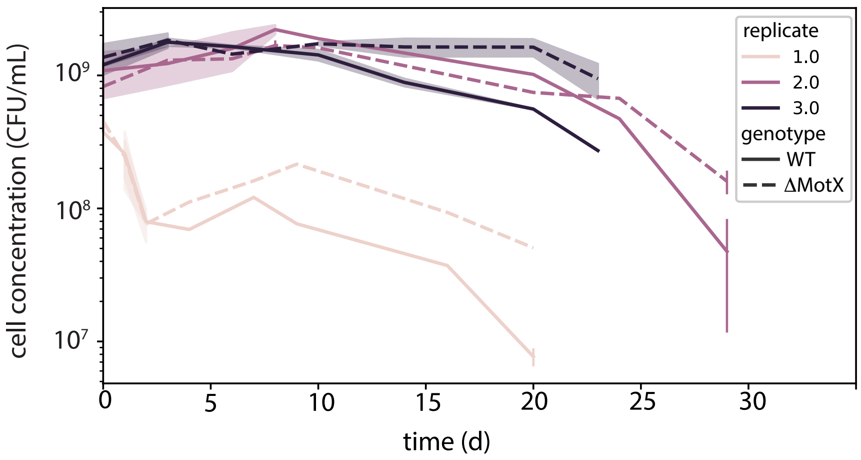

**Fig. S2. Dissipation of growing populations**. The raw measured heat flux Q*(t) and the heat flux per biomass Q(t), computed from an initial biomass of OD=0.002, a specific growth rate of 1.1 $h^{-1}$ and a carrier capacity of OD=0.32 as measured in experiments with the same calorimetry vials at 27C.

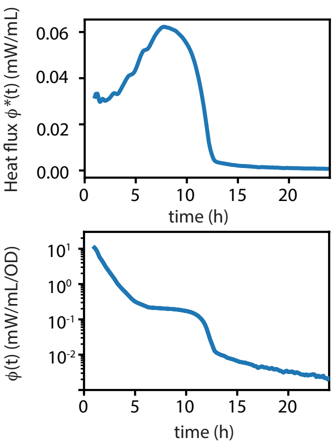

**Fig. S3. Alternative motility defect mutant in *E. coli***. Dissipation based on *E. coli* the difference between WT and △MotB (green). For comparison the dissipation based on the difference between WT and △MotA (black, see Fig. 4) is also shown.

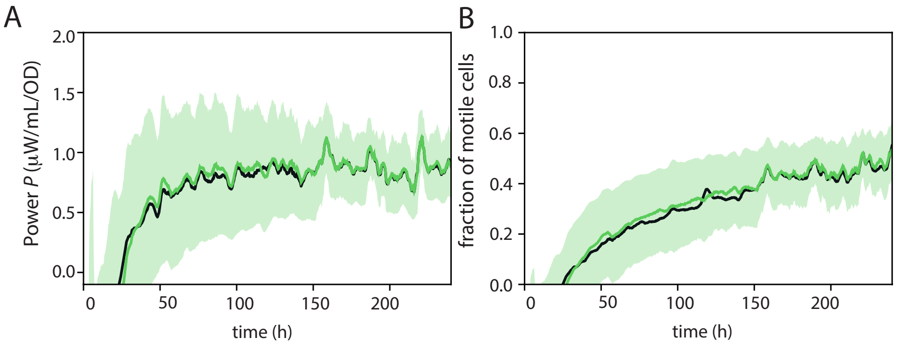

**Fig. S4. Comparison with RFT-Lighthill and Slender Body theory (SBT).** The dissipation per cell as measured with calorimetry (purple) and the theoretical estimates based on RFT using the parameters of Lighthill (red) and SBT (green). Shaded area represents 1 standard error of the mean based on 4 biological replicates, as in Fig. 3. For the theoretical estimates, squares and points represent respectively the mean and individual data points, with the error bars representing 1 s.e.m.

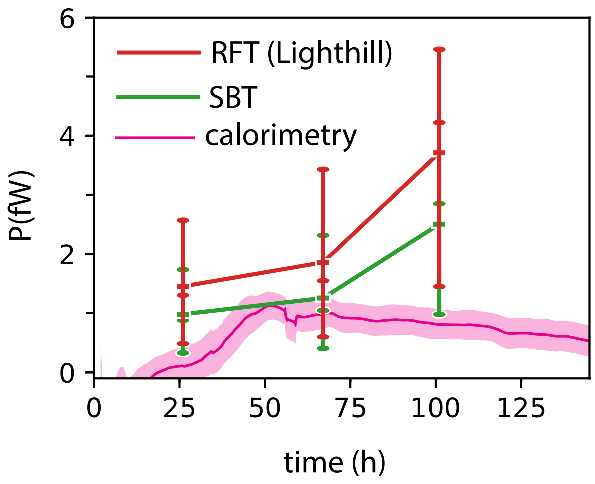

**Fig. S5. Swimming efficiency according to GH-RFT based on cell and flagellar morphology**. The different morphological parameters as illustrated were extracted from scanning-electron-microscopy images of *Vibrio coralliilyticus* starved for 24h (Supplementary Table 1). In all the panels, a dot represents a single cell and Pearson's correlation coefficient is computed for each parameter pair, as indicated in the plot by the blue line. The bottom row is the flagellar efficiency $\eta_f = \eta_{GH}$ ($\eta$ =1 corresponds to fully efficient).

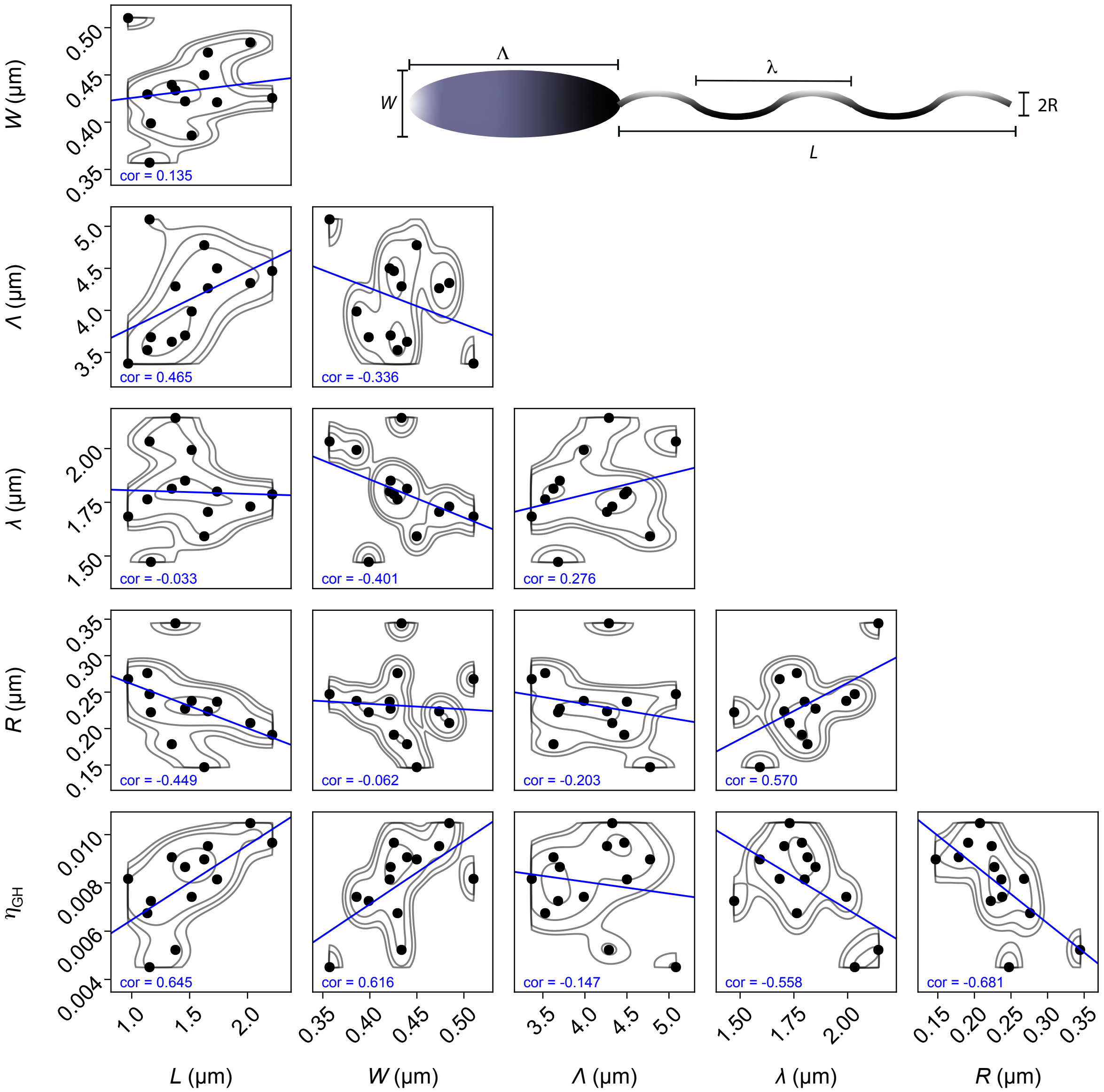

#### 3. Supplementary Tables

**Table S1. *Vibrio coralliilyticus* YB1 cell and flagella morphologies after 24 h of starvation as measured with SEM. (11). Each row represents the same individual cell. Bottom row is the average value of all cells.**

| cell length ($\mu$m) | cell width ($\mu$m) | flagellar length ($\mu$m) | pitch ($\mu$m) | wave amplitude ($\mu$m) |
|---|---|---|---|---|
| 1.1538 | 0.3571 | 5.0824 | 2.0330 | 0.4945 |
| 1.3462 | 0.4396 | 3.6264 | 1.8132 | 0.3571 |
| 1.5177 | 0.3859 | 3.9871 | 1.9936 | 0.4759 |
| 1.7368 | 0.4211 | 4.5000 | 1.8000 | 0.4737 |
| 1.6579 | 0.4737 | 4.2632 | 1.7053 | 0.4474 |
| 0.9694 | 0.5102 | 3.3673 | 1.6837 | 0.5357 |
| 1.3776 | 0.4337 | 4.2857 | 2.1429 | 0.6888 |
| 2.2128 | 0.4255 | 4.4681 | 1.7872 | 0.3830 |
| 2.0242 | 0.4844 | 4.3253 | 1.7301 | 0.4152 |
| 1.6263 | 0.4498 | 4.7751 | 1.5917 | 0.2941 |
| 1.4610 | 0.4221 | 3.7013 | 1.8506 | 0.4545 |
| 1.1350 | 0.4294 | 3.5276 | 1.7638 | 0.5521 |
| 1.1656 | 0.3988 | 3.6810 | 1.4724 | 0.4448 |
| 1.4911 | 0.4332 | 4.1223 | 1.7975 | 0.4628 |

**Table S2. Optical Density (OD) Measurements by Species, Mutation, and Condition**

| Species | Mutation | Condition | OD | Biological Replicate |
|---|---|---|---|---|
| *E. coli* | WT | MotM | 0.1540 | 1 |
| *E. coli* | MotB | MotM | 0.3100 | 1 |
| *E. coli* | MotA | MotM | 0.3200 | 1 |
| *E. coli* | WT | MotM | 0.2500 | 2 |
| *E. coli* | MotB | MotM | 0.2450 | 2 |
| *E. coli* | MotA | MotM | 0.2700 | 2 |
| *E. coli* | WT | MotM | 0.2350 | 3 |
| *E. coli* | MotB | MotM | 0.2550 | 3 |
| *E. coli* | MotA | MotM | 0.2750 | 3 |
| *V. corr* | WT | ASW | 0.6000 | 1 |
| *V. corr* | MotX | ASW | 0.3700 | 1 |
| *V. corr* | WT | ASW | 1.0400 | 2 |
| *V. corr* | MotX | ASW | 1.0800 | 2 |
| *V. corr* | WT | ASW | 0.3200 | 3 |
| *V. corr* | MotX | ASW | 0.4000 | 3 |
| *V. corr* | WT | ASW | 0.3100 | 4 |
| *V. corr* | MotX | ASW | 0.3100 | 4 |
| *V. corr* | WT | LowS | 0.7519 | 1 |
| *V. corr* | WT | LowS | 0.7282 | 2 |
| *V. corr* | WT | LowS | 0.7576 | 3 |
| *V. corr* | MotX | LowS | 1.2791 | 1 |
| *V. corr* | MotX | LowS | 0.9799 | 2 |
| *V. corr* | MotX | LowS | 1.0608 | 3 |
| *V. corr* | WT | MedS | 0.4769 | 1 |
| *V. corr* | WT | MedS | 0.5399 | 2 |
| *V. corr* | WT | MedS | 0.4087 | 3 |
| *V. corr* | MotX | MedS | 1.3044 | 1 |
| *V. corr* | MotX | MedS | 1.0397 | 2 |
| *V. corr* | MotX | MedS | 1.0422 | 3 |
| *V. corr* | WT | HighS | 0.5954 | 1 |
| *V. corr* | WT | HighS | 0.6822 | 2 |
| *V. corr* | WT | HighS | 0.6694 | 3 |
| *V. corr* | MotX | HighS | 1.3204 | 1 |
| *V. corr* | MotX | HighS | 1.0377 | 2 |
| *V. corr* | MotX | HighS | 1.1201 | 3 |

**Movie S1. Representative video of *Vibrio coralliilyticus* YB1 WT. Cells were grown in marine broth and washed in artificial seawater, and observed using phase contrast microscopy (10X) at 30 frames per second. As background subtraction, the median pixel value of each pixel over the time series duration (900 frames) is subtracted. For clarity, the an area of 512 by 512 pixels (original image size 2044x20248 pixels) is displayed.**

**Movie S2. Representative video of *Vibrio coralliilyticus* YB1 ΔMotX under the same conditions as WT in Supplementary video 1**
